# Prestimulus Activity in the Cingulo-Opercular Network Predicts Memory for Naturalistic Episodic Experience

**DOI:** 10.1101/176057

**Authors:** Noga Cohen, Aya Ben-Yakov, Jochen Weber, Micah G. Edelson, Rony Paz, Yadin Dudai

## Abstract

Human memory is strongly influenced by brain states occurring before an event, yet we know little about the underlying mechanisms. We found that activity in the cingulo-opercular network (including bilateral anterior insula and anterior prefrontal cortex) seconds before an event begins can predict whether this event will subsequently be remembered. We then tested how activity in the cingulo-opercular network shapes memory performance. Our findings indicate that prestimulus cingulo-opercular activity affects memory performance by opposingly modulating subsequent activity in two sets of regions previously linked to encoding and retrieval of episodic information. Specifically, higher prestimulus cingulo-opercular activity was associated with a subsequent increase in activity in temporal regions previously linked to encoding and with a subsequent reduction in activity within a set of regions thought to play a role in retrieval and self-referential processing. Together, these findings suggest that prestimulus attentional states modulate memory for real-life events by enhancing encoding and possibly by dampening interference from competing memory substrates.

---

Successful memory formation is associated with enhanced activity in brain regions linked to encoding such as the fusiform and medial temporal regions (Kim 2011) and with reduced activity in regions associated with retrieval and self-referential processes, such as the precuneus and posterior cingulate cortex (Kim et al. 2010). Thus far, the examination of the neural correlates of memory formation has focused mainly on the brain activity occurring during (e.g., Davachi et al. 2003; Eichenbaum et al. 2007; Kim et al. 2010; Kim 2011) or following (Tambini et al. 2010; Ben-Yakov and Dudai 2011; Staresina et al. 2013; Ben-Yakov et al. 2014; Tompary et al. 2015) the presentation of the memoranda. Processes occurring before the onset of an event, however, can also shape memory formation. While some studies probed the prestimulus brain activity that predicts memory performance (for review see Cohen et al. 2015), it is yet unclear how prestimulus activity and activity during the stimulus interact to modulate encoding.

Prior studies that have examined memory-predictive prestimulus activity found that activity in regions such as the hippocampus, amygdala, and midbrain (Adcock et al. 2006; Addante et al. 2015; de Chastelaine and Rugg 2015; Mackiewicz et al. 2006; Park and Rugg 2010; Sadeh et al. 2019; Wittmann et al. 2007) predicted whether an upcoming event will be later remembered or forgotten. Specifically, differences between subsequently remembered and subsequently forgotten stimuli were observed several seconds prior to stimulus onset. It has been suggested that prestimulus activity in these regions enhances memory formation by preparing the system to encode the upcoming stimulus (e.g., by lowering the threshold for LTP in the medial temporal lobe; Frey et al. 1993; Huang and Kandel 1995; Otmakhova and Lisman 1996). Electrophysiological data suggest that enhanced pre-stimulus theta and alpha activity predict subsequent memory success (Fell et al. 2011; Koen et al. 2018; Gilboa and Moscovitch 2017; Guderian et al. 2009; Merkow et al. 2014; Strunk and Duarte 2019; Sweeney-Reed et al. 2016), possibly via involvement of frontal (Gilboa and Moscovitch 2017; Koen et al. 2018; Otten et al. 2006) as well as temporal (Fell et al. 2011; Guderian et al. 2009; Merkow et al. 2014), and thalamic (Sweeney-Reed et al. 2016) regions.

While the aforementioned studies provide important insights regarding the prestimulus brain correlates of memory formation, their findings may have been affected by specific task characteristics. Specifically, in most of these studies a cue informed the participant of the content of the upcoming to-be-remembered target. For example, memory-predictive prestimulus activity in the amygdala was found following a cue predicting a subsequent appearance of an unpleasant picture (Mackiewicz et al. 2006), while memory-predictive prestimulus activity in the midbrain was found following a cue predicting a rewarding target (Adcock et al. 2006). In addition, these studies tested episodic memory for static stimuli, such as words or pictures, and it is unclear whether the same brain circuits are involved in the encoding of dynamic, naturalistic events. Furthermore, these studies did not examine the link between prestimulus activity and online stimulus activity and thus provide only indirect evidence as to how prestimulus activity modulates memory performance. The aim of the current study was therefore twofold: 1) identify prestimulus activity that predicts memory outcome in naturalistic settings, and 2) offer a mechanistic account for the role of this activity in shaping memory formation.

We first identified memory-predictive prestimulus activity using a subsequent memory functional magnetic resonance imaging (fMRI) study (Experiment 1). Participants were presented with naturalistic memoranda (brief narrative movie clips) and their memory for the main occurrence (gist) in each of the clips was tested following the scan using a cued-recall task. After identifying memory-predictive prestimulus activity, we tested how it may shape memory performance by modulation of intra-stimulus activity. Specifically, following the findings showing memory-predictive prestimulus activity in the cingulo-opercular network, which is commonly considered to be associated with top-down control of attention, we conducted two multi-level mediation analyses to test the following predictions regarding prestimulus cingulo-opercular activity: 1) it enhances memory performance by boosting online encoding activity; 2) it enhances memory by suppressing task-unrelated, self-generated thoughts. We then tested whether findings of Experiment 1 are replicated in an independent data-set (Experiment 2) in which subsequent memory was tested one day following the scan, using an a-priori dedicated mask of the cingulo-opercular network.

## Experiment 1

Experiment 1 included a subsequent memory task in which participants were presented with naturalistic memoranda (brief narrative movie clips) in an fMRI scanner. Memory for the clips was tested outside the MRI about 20 minutes following the scan. In addition to BOLD signal, we collected eye tracking data (eye-movements, blinks and pupil size). These measures were used to control for participants’ engagement and arousal during the task and are reported in the Supplementary Information.

## Materials and Methods

### Participants

Experiment 1 included 28 participants (12 male, mean age = 25.5 ± 3.2). Two participants were excluded due to excessive head movements, three participants were excluded due to technical problems during the Study session, one participant was excluded due to low memory performance (correctly recalled less than 10% of the movies; 14 trials), hence the resulting sample included 22 participants (9 males, mean age = 25.7 ± 3.4). The study was approved by the ethics committee of the Weizmann Institute of Science, and all subjects gave informed consent prior to the experiment.

### Stimuli

Each participant viewed 160 audiovisual clips (Ben-Yakov and Dudai 2011a). Of these clips, 140 were narrative movie clips and were used for the current analysis. For other purposes, beyond the scope of the current paper, the clips were preceded by either a “remember” or a “look” instruction cue. Participants were told that the memory test following the scan will include only clips that are preceded by a “remember” cue, and they will not be tested on clips that follow a “look” cue. Half of the trials included a “remember” cue while the other half included a “look” cue. The cue appeared on the screen for 7/9/11 s and was followed by a clip that lasted 8 s. Memory was subsequently tested for all clips.

### Experimental protocol

#### Study session

The Study session took place in an fMRI scanner and was divided into four scanning runs. Each run started and ended with the presentation of a blank screen for 10 s. Each trial (see Figure 1 for an example) started with a fixation cross for 2 s. Then, an instruction word (“remember”/”look”) was presented for a jittered length (7/9/11 s with an average of s). In order to eliminate temporal anticipation effects, the distribution of instruction lengths was determined using the nonaging foreperiod distribution (Niemi and Näätänen 1981). Specifically, there was a 50 % probability that the clip will appear in any given foreperiod. This structure was designed specifically to eliminate participants’ ability to predict when the clip will appear following the onset of the instruction cue. There were 80 trials in the 7 s foreperiod, 40 trials in the 9 s foreperiod, and 20 trials in the 11s foreperiod. In addition, 20 catch trials were included, in which the clip was a visually scrambled clip accompanied by a non-distinctive background noise. Catch trials were always preceded by the longest foreperiod duration (11 s instruction), thus making it impossible for participants to predict whether the instruction would be followed by a narrative or by a control clip. Following the instruction word, a clip was presented for 8 s. Each trial ended with a fixation cross for 3 s.

**Figure 1.**
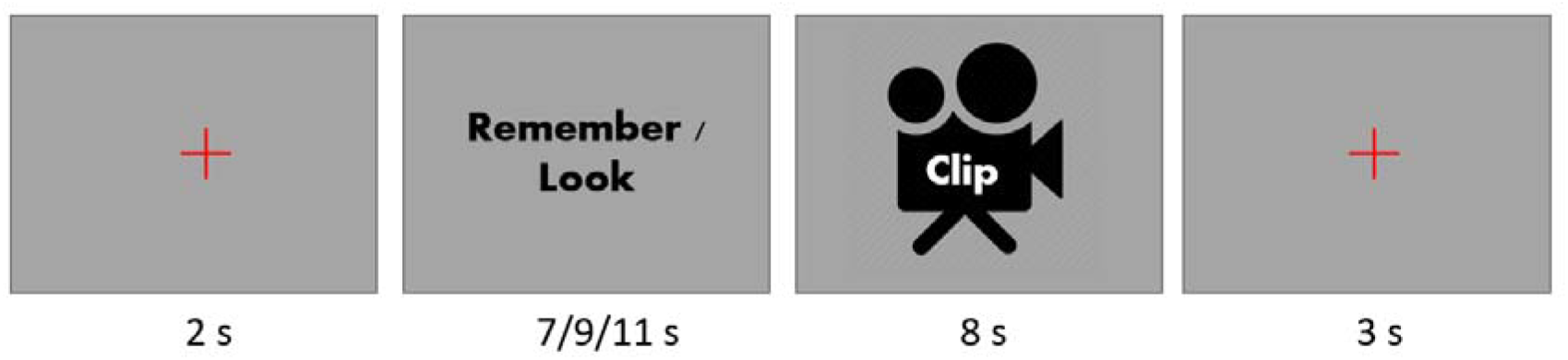
Example of a trial in the Study session, which was conducted while participants undergo fMRI scanning. On each trial, participants received an instruction cue (“remember” or “look”) prior to a brief (8 seconds) movie clip. They were told that following the scan, they would perform a memory test asking them about the gist of the clips that follow a “remember” instruction.

#### Test session

The Test session took place outside the scanner, 20 min after the Study session. Participants were informed beforehand about the format of the Test session. The Test session consisted of a cued recall test; for each of the clips, one question about the main gist of the clip (e.g., “What did the parents say to their son?”) appeared bellow a representative frame from that clip. Clips that received a correct answer were labeled as “remembered” and clips that received no answer or a wrong answer were labeled as “forgotten”. Participants were instructed to provide a response only if they remembered the main gist of the clip, hence most (mean = 91%, SD = 7.5) of the “forgotten” labeled clips were those in which the participant did not provide a response (i.e., only 9% of the forgotten clips were clips with an erroneous response). In cases where it was not completely clear whether an answer was correct, the corresponding clip was labeled as “unclear” and excluded from analysis (mean = 5.5%, SD = 2.67). The test probed memory for all clips, including those preceded by the “look” instruction. The reliability of the scoring was verified by two additional raters (see Supplementary Information).

### fMRI acquisition and data analysis

The experiment was carried out on a Siemens 3T Trio Magnetom scanner at the Weizmann Institute of Science, Rehovot, Israel. BOLD contrast was obtained using a gradient-echo EPI sequence (FOV – 216 cm, matrix size – 72 x 72, voxel size – 3 x 3 x 4 mm³, TR/TE/FA = 2,000 ms / 30 ms / 75 degrees, 32 axial slices). A T1-weighted 3D MPRAGE sequence was used to collect anatomical scans (voxel size – 1 x 1 x 1 mm³, TR/TE/FA = 2,300 ms / 2.98 ms / 9 degrees).

### fMRI data pre-processing

fMRI data were processed and analyzed using Statistical Parametric Mapping software (SPM8; Wellcome Department of Imaging Neuroscience, London, UK) with MATLAB 7.14.0 (the Mathworks, USA). Pre-processing included slice timing correction to the middle slice, motion correction using realignment to the first volume, and co-registration to the individual high-resolution anatomical image. Then, normalization to Montreal Neurological Institute (MNI) space (Mazziotta et al. 1995) was performed using the unified segmentation approach (Ashburner and Friston 2005). Images were then spatially smoothed with a 6-mm full width at half maximum (FWHM) Gaussian kernel. Voxel size following pre-processing was set to be 3 x 3 x 3 mm³.

### fMRI data analysis

Prestimulus activity during the instruction time-window was modeled using box-car epochs with variable durations (i.e., from instruction onset to clip onset, lasting 7, 9, or 11 seconds), convolved with the canonical hemodynamic response function (HRF). For each participant, a set of eight regressors were constructed, for all possible combinations of instruction type (remember/look) and clip type (remembered/forgotten/control/x). This resulted in the following conditions: remember-remembered, remember-forgotten, remember-control, remember-X, look-remembered, look-forgotten, look-control, look-X. In addition, six motion realignment nuisance regressors, as well as white matter (WM) and cerebrospinal fluid (CSF) regressors, were added to the GLM, and a high-pass filter of 100 s was applied. The BOLD signal of the WM and CSF were used to exclude signals in areas of no interest, such as white matter and ventricles (Behzadi et al. 2007; Chai et al. 2012). These regressors were calculated by averaging the BOLD signal in WM and CSF masks defined by the segmentation function of SPM8. Z-score values of these BOLD signals were entered as covariates of no interest to the first-level models to account for global variability unrelated to the task (Power et al. 2017). First level models were computed using either (1) prestimulus phase regressor, (2) prestimulus phase in addition to clip phase regressors. The differences observed between (1) and (2) were small and therefore we chose to focus on the more parsimonious model (1) and present model (2) in the supplementary information. The single-subject contrasts were then taken to a standard full factorial ANOVA with the relevant task conditions as factors (remember-remembered, look-remembered, remember-forgotten, look-forgotten). We used a second-level contrast to assess the main effect of interest (remembered > forgotten, Figure 2a). See Supplementary Information for the main effect of instruction type (remember > look) and for control analyses showing no indication for sequential effects or modulation of the main effect by instruction duration or by prestimulus arousal (indicated by pupil size). The interaction between instruction type and memory performance revealed some differences is some of the regions (see Supplementary Information for time courses representing all four conditions; SI4) but these differences did not reach significance (p < 0.001, cluster pFWE < 0.05) and therefore we collapsed across the two instruction types (Figure 2b).

**Figure 2.**
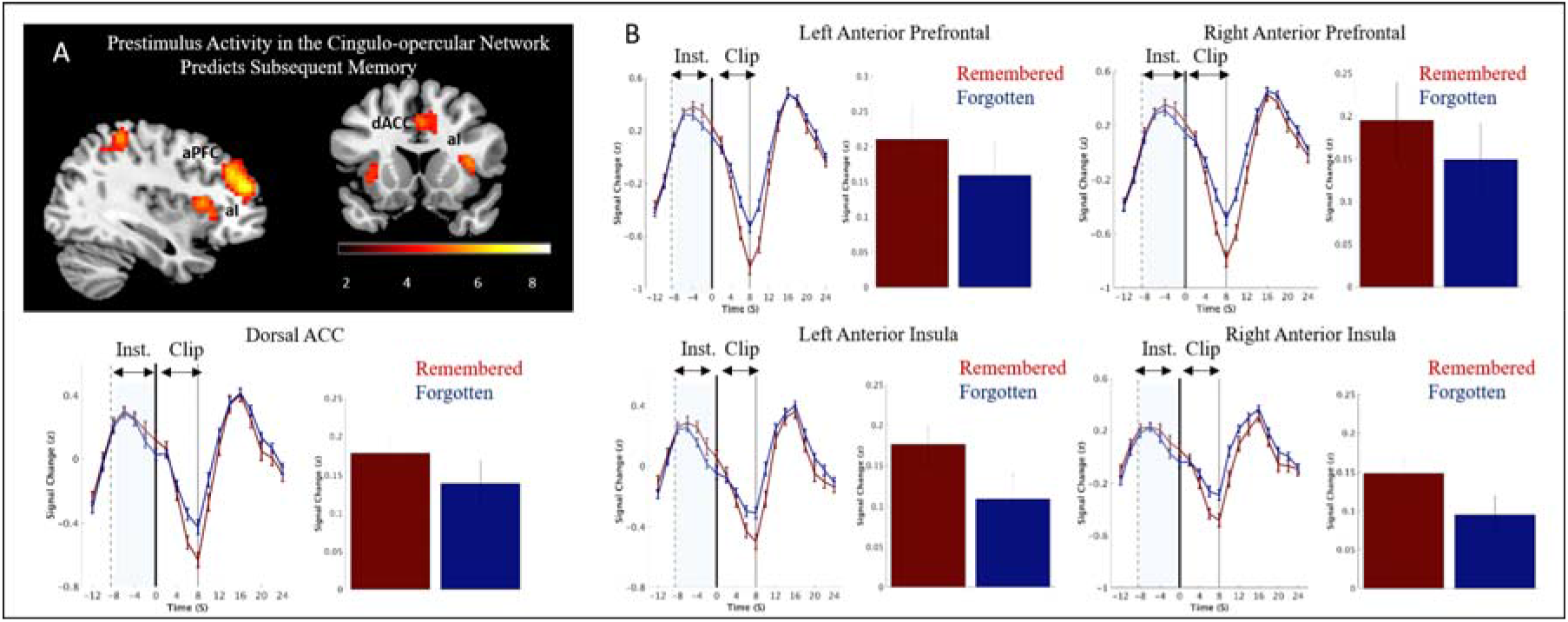
**A)** Regions demonstrating higher prestimulus BOLD activity for remembered vs. forgotten clips (*p* < 0.001, cluster pFWE < .05) in Experiment 1. Data are shown on sagittal and axial slices of an MNI template. aI-anterior insula; aPFC - anterior prefrontal; dACC - dorsal anterior cingulate cortex. **B)** For illustration purposes, mean group BOLD signal (after z scoring each time course) for remembered and forgotten clips in Experiment 1 were extracted from regions of the cingulo-opercular network using a functional ROI. The solid black lines indicate the onset of clip presentation, the solid gray lines indicate the offset of the current clips, and the dashed lines represent the mean onset of the instruction cue. Error bars represent the standard error of the mean across subjects (random effects). The bar figures represent the mean activity during the instruction time-window for each of the conditions.

For the whole-brain analysis, we used a voxel-level threshold of p < 0.001 and a cluster-level threshold of pFWE < 0.05, using SPM’s built-in Gaussian Random Fields (GRF) correction procedure. The cluster-forming threshold (CFT, p < 0.001) was chosen to approximately correctly account for the expected false-positive rate using GRF (Eklund et al. 2016). For illustration purposes, time courses were extracted by Z-scoring the raw BOLD signal for each run of each participant. The time courses were then averaged across all events from the same type (remembered, forgotten) within each participant and then across participants.

### Linking prestimulus and online activity

We sought to reveal whether the link between prestimulus activity and memory performance was mediated by online stimulus activity. This consisted of two phases: 1) Identifying regions in which online (intra-stimulus activity) was correlated with the prestimulus activity in the cingulo-opercular regions, and 2) running a mediation analysis to determine whether online activity mediated the link between prestimulus activity and memory.

#### 1) Identifying mediators

A parametric analysis was conducted to identify regions in which online activity was correlated with prestimulus cingulo-opercular activity. We created parametric weights from the per-trial prestimulus response in cingulo-opercular regions, defined by the whole-brain analysis described above for the remembered > forgotten contrast (averaging over all regions identified by the contrast as they were highly correlated). We then ran a GLM with a regressor for online activity, using the prestimulus weights for parametric modulation. This yielded regions whose activity during the stimulus was either correlated or anti-correlated with prestimulus cingulo-opercular activity. Since our interest was to test whether the link between prestimulus activity and subsequent memory performance is mediated by online activity, the mediation analysis required collapsing across the remembered and forgotten trials (to avoid a dependency between the parametric modulator and the main effect of memory performance). Therefore, the model included two regressors (narrative clips and control clips), and their parametric modulation regressors. A second-level analysis (one sample t-test, voxel-level threshold of p < 0.001 and a cluster-level threshold of pFWE < 0.05) was conducted only on the parametric regressor of the narrative clip events. We computed both the positive (1 coded) and negative (−1 coded) contrasts for the parametric regressor.

#### 2) Mediation model

The aim of the second phase was to determine whether the online activity identified in phase 1 mediated the link between prestimulus cingulo-opercular activity and memory. For this purpose, two additional regression models were estimated, in which we constructed separate regressors for each trial (e.g., Rissman et al. 2004). In the first of these single-trial models, we modeled the prestimulus phase of each event (total of 160 regressors), convolved with the canonical HRF. The second model was constructed analogously, but the actual clip stimulus period was modeled (into an equal number of 160 regressors). For each of the prestimulus and clip stimulus periods, we then extracted the ROI-averaged beta (amplitude) estimates for the prestimulus whole-brain remembered > forgotten contrast, and for two sets of regions that were identified using the parametric model described above. The prestimulus betas (from the cingulo-opercular ROI) and the stimulus betas (from the parametric model) were then, together with memory performance (coded as 0 for forgotten and 1 for remembered), subjected to two Bayesian multi-level logistical mediation analyses, one for the positive regions and one for the negative ones. The mediation analyses were conducted using the bmlm R package (Vuorre and Bolger 2017), which uses the RStan interface to conduct the Bayesian inference (Stan Development Team, 2016). For the regression parameters, the prior distributions are zero-centered Gaussians, with user-defined standard deviations (defaults to 1000). For each path (a, b, c, c’, ab) we present the fixed-effect parameter, and its associated credible intervals (95% mass of the marginal posterior distribution).

## Results

### Memory performance

Participants remembered 41% ±3% of the clips. We tested to what extent memory performance differed between the two instruction types (“remember”/”look”) by comparing the percent of remembered items (remembered/(remembered+forgotten)) between the two instruction types. A paired t-test revealed an effect for instruction type, *t* (21) = 5.39, p < 0.0001, with a higher percent of remembered clips in “remember” vs “look” trials (mean number of events in each trial type; remember-remembered: 32 clips, look-remembered: 23 clips, remember-forgotten: 34 clips, look-forgotten: 44 clips).

### Prestimulus activity in the cingulo-opercular network predicts subsequent memory

In order to identify regions demonstrating higher prestimulus activity for subsequently remembered vs. forgotten clips, we conducted a whole-brain analysis (cluster-forming threshold *p* < 0.001, cluster-pFWE < 0.05), in the instruction time window. This analysis yielded significant activity in a set of regions usually considered to be part of the cingulo-opercular network, as well as two regions (left precuneus and left inferior parietal) sometimes considered to be part of the frontoparietal network (see Figure 2a and Supplementary Information Table SI2). These regions were also found to be co-activated with the cingulo-opercular network during the initiation of task goals (Dosenbach et al., 2006). As results indicated robust involvement of all cingulo-opercular regions, we decided to focus on this network in all further analyses. Illustration of this effect can be seen in Figure 2b, which depicts the mean BOLD signal for remembered and forgotten clips, extracted from regions of the cingulo-opercular network (see Supplementary Information SI4 for the time-course of the response to remembered/forgotten events divided by instruction).

As differences due to instruction type are not the main focus of the current work, we present these findings in the supplementary information. The contrast “remember” vs “look” (*p* < 0.001, cluster-pFWE < 0.05) revealed bilateral occipital activity (see Supplementary Information SI2), and the interaction between instruction type (“remember”/”look”) and memory (remembered/forgotten) did not reveal significant activations (*p* < 0.001, cluster-pFWE < 0.05).

### Linking prestimulus activity to online activity

The cingulo-opercular network is usually associated with adaptive control of attention (Dosenbach et al. 2008) and therefore we predicted that this network may set the stage for encoding by shaping online clip-related activity. In order to test this prediction we 1) probed for possible mediators linking prestimulus cingulo-opercular activity and memory performance using a parametric analysis; and 2) conducted two multi-level logistic Bayesian mediation models to examine the role of these potential mediators in the association between cingulo-opercular activity and memory performance.

#### 1) Cingulo-opercular activity predicts online activity

A parametric analysis was conducted in order to explore whether prestimulus activity in the cingulo-opercular network modulated activity during clip presentation. A whole-brain analysis was used to detect brain regions that, during the clip time-window, were positively or negatively associated with cingulo-opercular prestimulus activity. Regions that were positively associated with prestimulus activity were regions showing an increase in their activity during clip presentation following higher prestimulus cingulo-opercular activity. Regions that were negatively associated with prestimulus activity were regions showing a *reduction* in their activity during clip presentation following higher cingulo-opercular activity in the prestimulus phase.

The whole-brain analysis (see Supplementary Information SI9) for positive modulation by cingulo-opercular activity revealed significant activations in the fusiform gyrus and middle temporal regions. The whole-brain analysis for the negative parametric modulation showed significant activations in a set of regions that included the posterior cingulate, precuneus, and striatum.

#### 2) Online activity mediates the cingulo-opercular – memory link

Two mediation analyses were conducted to determine whether the link between prestimulus cingulo-opercular activity and memory was mediated by activity in the set of brain regions found in the parametric analysis. Specifically, we tested whether activity in regions found in the positive parametric contrast and activity in regions found in the negative parametric contrast mediate the cingulo-opercular - memory link. Both analyses indicated partial mediation, suggesting that more than 40% of the link between prestimulus cingulo-opercular activity and successful memory performance was mediated by an increase in clip-related activity in a set of temporal regions (Figure 3A) and by a decrease in clip-related activity in a set of regions that included the precuneus, striatum, and cingulate cortex (Figure 3B). Specifically, measuring the path coefficients for a standard three-variable path model that used activity from regions revealed in the positive parametric contrast (+1) as a mediator demonstrated credible relationships between prestimulus cingulo-opercular activity and memory performance (path c: b = 1.12, [0.83, 1.41]), prestimulus cingulo-opercular activity and activity in regions found in the positive parametric contrast (path a: b = 0.2 [0.18, 0.23]), and between activity in regions found in the positive parametric contrast and memory performance (path b: b = 2.19 [1.68, 2.71]). Furthermore, the relationship between cingulo-opercular prestimulus activity and memory was reduced when activity in regions found in the positive parametric contrast was included in the model (path ab: b = 0.45 [0.33, 0.58]), although the relationship between the prestimulus cingulo-opercular activity and memory was still present (path c’: b = 0.67 [0.39, 0.94]).

**Figure 3.**
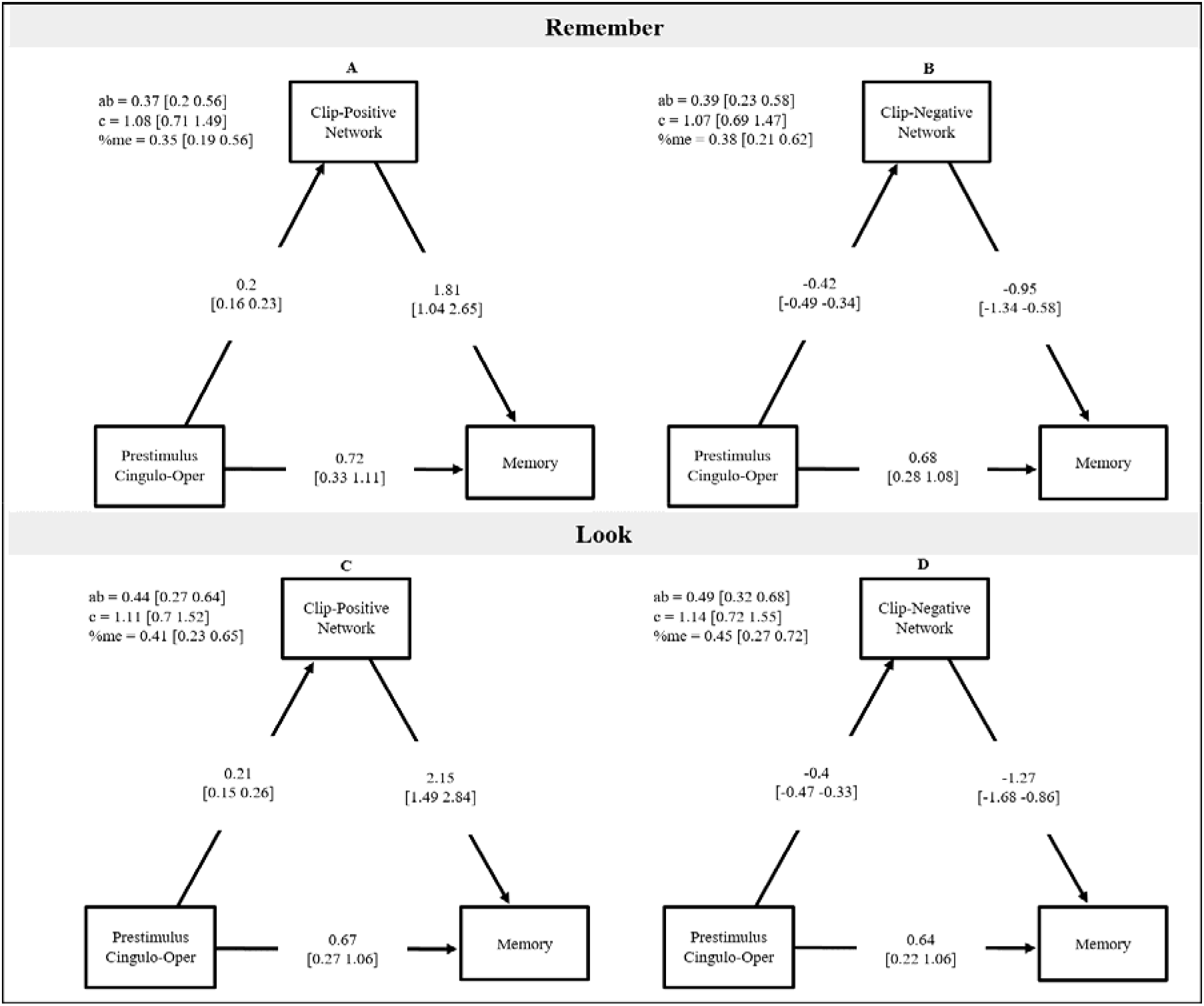
Multi-level mediation analyses assessing the contribution of regions implicated in the parametric analysis for the two experiments. A, C) A model including activity from the set of regions found in the positive contrast of the parametric analysis (Clip-Positive Network) as a mediator. B, D) A model including activity from the set of regions found in the negative contrast of the parametric analysis (Clip-Negative Network) as a mediator.

Measuring the path coefficients for the model that used activity from regions revealed in the negative parametric contrast (−1) as mediator revealed credible relationships between prestimulus cingulo-opercular activity and memory performance (path c: b = 1.12, [0.82, 1.4]), prestimulus cingulo-opercular activity and activity revealed in the negative parametric contrast (path a: b = −0.41 [−0.47 −0.35]), and between activity revealed in the negative parametric contrast and memory performance (path b: b = −1.19 [−1.47, −-0.92]). As in the aforementioned model, the relationship between prestimulus cingulo-opercular activity and memory was reduced when the mediator (activity in regions found in the negative parametric contrast) was included in the model (path ab: b = 0.48 [0.36, 0.62]), although the relationship between the prestimulus cingulo-opercular activity and memory was still present (path c’: b = 0.63 [0.34, 0.92]).

Thus, the statistical criteria for partial mediation were met in both models, indicating that the link between memory performance and prestimulus activity in the cingulo-opercular network was partially accounted for by the increased online activity in a network including temporal regions and decreased online activity in a network including the posterior precuneus, cingulate and striatum.

Importantly, testing these mediations separately for the “remember” and “look” instruction type did not reveal a difference between these two types of trials. Namely, the mediation was present for both instruction types (see SI10). This suggests that the mediating role of online activity in the cingulo-opercular – memory link is not dependent on task-goals and is present also when there is no intention to encode.

## Experiment 2

Data from Experiment 2 were used to replicate the findings of Experiment 1, as well as to generalize the findings to naturalistic situations. Therefore, in contrast to Experiment 1, Experiment 2 did not include an instruction cue prior to the clip. In addition, subsequent memory was assessed a day after the scan, making it possible to see whether the effects hold when the recall is not assessed immediately after encoding. Furthermore, in order to confirm the role of the cingulo-opercular network in the effects observed, we probed this network using a dedicated mask.

## Materials and methods

A data-set from an independent study previously conducted in our lab (Experiment 3 in (Ben-Yakov and Dudai 2011a) was used in the current experiment.

### Participants

Experiment 2 included 21 participants. Three participants were excluded due to low memory performance (correctly recalled less than 10% of the movies; 12 trials), hence the resultant sample included 18 participants (11 males, mean age = 26.7 ± 2.8). The study was approved by the ethics committee of the Weizmann Institute of Science, and all subjects gave informed consent prior to the experiment.

### Stimuli

Each participant viewed 128 clips. Of these clips, 112 were narrative movie clips and were used in the current analysis. An additional 16 uneventful clips were modeled as nuisance regressors as this condition was outside the scope of the current paper. The clips were of varied lengths (32 clips of 8 s, 64 clips of 12 s, and 16 clips of 16 s). The design also included 4 brief blocks of a go/no-go task, aimed at increasing participants’ alertness (Ben-Yakov et al., 2011).

### Experimental protocol

#### Study session

The Study session took place in an fMRI scanner and was divided into two scanning runs. The clips were presented in random order; each clip was preceded by a fixation screen of jittered length (8/10/12/14/16 s with an average of 10.75 s).

#### Test session

The Test session was similar to the one used in Experiments 1, but was administered one day following the Study session. As in Experiment 1, clips were coded as “remembered”, “forgotten”, or “unclear” (mean = 4.5%, SD = 2.09). Of the forgotten clips, 89% (SD = 23) were clips in which the participant did not provide a response (mean number of events in each trial type; remembered: 30 clips, forgotten: 77 clips). As in Experiment 1, The reliability of the scoring was verified by two additional raters (see Supplementary Information).

### fMRI acquisition and data analysis

The experiment was carried out on a Siemens 3T Trio Magnetom scanner at the Weizmann Institute of Science, Rehovot, Israel. BOLD contrast was obtained using a gradient-echo EPI sequence (FOV – 24 cm, matrix size – 80 x 80, voxel size – 3 x 3 x 4 mm³, TR/TE/FA = 2,000 ms / 30 ms / 75 degrees, 36 axial slices). A T1-weighted 3D MPRAGE sequence was used to collect anatomical scans (voxel size – 1 x 1 x 1 mm³, TR/TE/FA = 2,300 ms / 2.98 ms / 9 degrees).

### Data pre-processing

See Experiment 1. In the current experiment, we omitted the first 15 volumes (during this time there was an audiovisual clip for accommodation to fMRI).

### Network definition

In order to replicate the findings of Experiment 1 and examine their specificity to the cingulo-opercular network, we created an a-priori mask of the cingulo-opercular network based on a study by Dosenbach et al (2007), using WFUpickatlas toolbox (Maldjian et al. 2003; http://fmri.wfubmc.edu/software/PickAtlas). As in Dosenbach et al’s paper, this mask included 12mm spheres around peak coordinates (see Table 2) of the right and left anterior insula (aI), right and left anterior prefrontal (aPFC), and dorsal anterior cingulate cortex (dACC).

### Data analysis

As in Experiment 1, prestimulus activity was modeled using box-car epochs convolved with the canonical hemodynamic response function (HRF) on the prestimulus time-window (8-16 sec preceding clip onset). For each participant, a set of five regressors were constructed, coding for the different prestimulus events (remembered/forgotten/control/x/go-nogo). In addition, six motion realignment nuisance regressors, as well as WM and CSF regressors, were added to the GLM, and a high-pass filter of 100 s was applied. As in Experiment 1, we computed the statistics for two additional control models (see Supplementary Information). The single-subject contrasts were then taken to a repeated-measures ANOVA with all task conditions as factors. A specific contrast assessed the main effect of interest (remembered > forgotten). A small volume correction (SVC) analysis using a threshold of pFWE < .05 (Friston et al. 1996) was performed on the cingulo-opercular mask.

### Linking prestimulus on online activity

Similar to Experiment 1, a parametric analysis was conducted to explore the role of prestimulus activity in shaping online clip-related activity. Then, multi-level logistic mediation analyses were conducted to examine whether the link between the observed prestimulus cingulo-opercular activity and subsequent memory is mediated by clip-related activity in candidate regions found in the parametric analysis. These analyses were identical to those used in Experiment 1, only that the prestimulus activity was defined by an a-priori mask of the cingulo-opercular network.

## Results

### Memory performance

Participants remembered 27.6±3.9% of the clips. The memory performance in Experiment 2, in which the recall test was administered one day after the study session, was lower than in Experiment 1 (41%), in which the recall test was 20 min after the termination of the study session.

### Prestimulus activity in the cingulo-opercular network predicts subsequent memory

A mask of the cingulo-opercular network was used in a small volume correction (SVC) analysis. This analysis revealed significant activations in all regions of the network (see Supplementary Information Table SI12 and Figure 4; results of a whole-brain analysis are presented in the Supplementary Information SI13).

**Figure 4.**
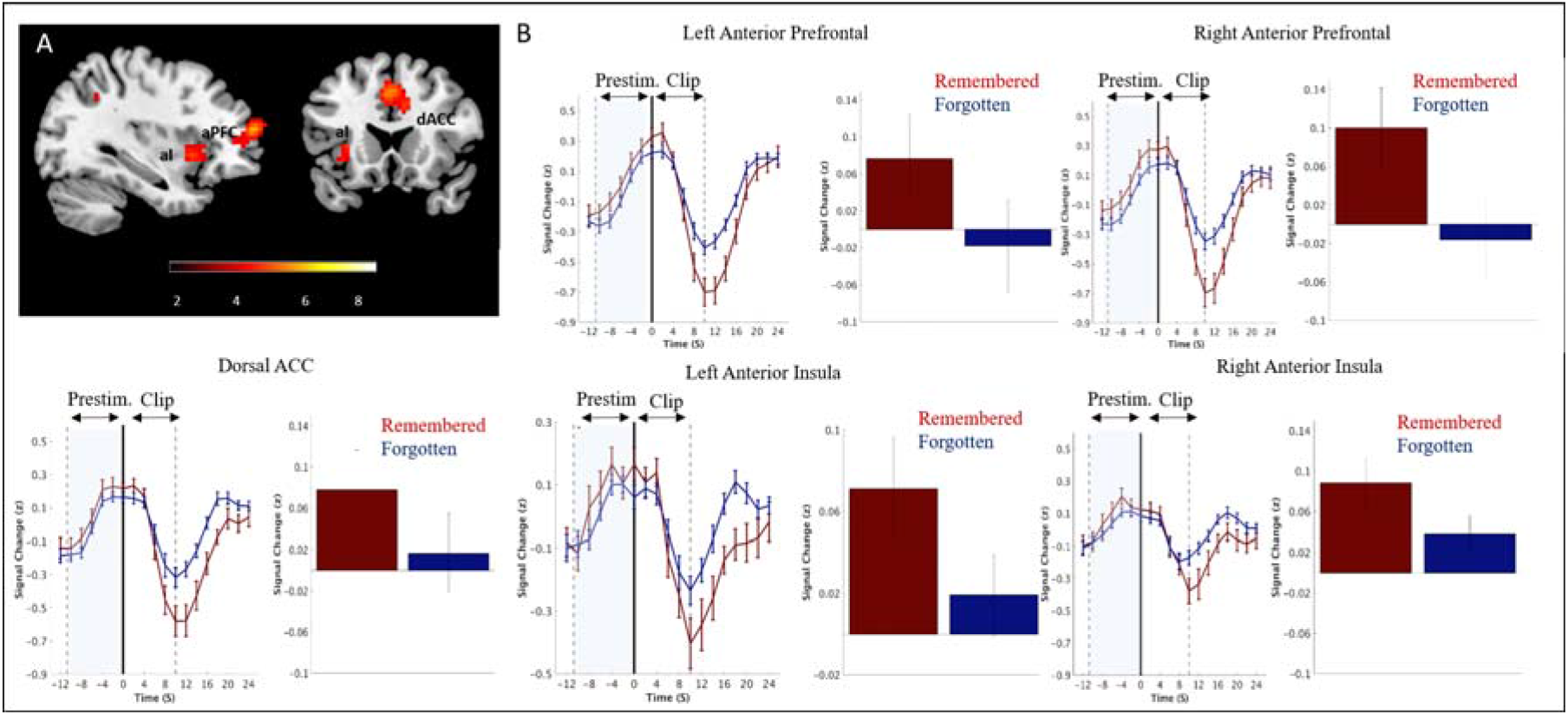
For illustration purposes, we present the mean group BOLD signal (after z scoring each time course) for remembered and forgotten clips in Experiment 2. BOLD activity was extracted from two example regions of the cingulo-opercular network using a functional ROI. The solid black lines indicate the onset of clip presentation, the solid gray lines indicate the offset of the current clips, and the dashed lines represent the mean onset of the instruction cue. Error bars represent the standard error of the mean across subjects (random effects). The bar figures represent the mean activity during the instruction time-window for each of the conditions.

### Linking prestimulus and online activity

#### 1) Online activity is correlated with prestimulus activity

A parametric analysis was conducted in order to explore whether prestimulus activity in the cingulo-opercular network modulated activity during clip presentation. Results replicated findings of Experiment 1; the whole-brain analysis (see Supplementary Information Table SI16) for positive modulation by cingulo-opercular activity revealed significant activations in a set of regions that included the fusiform gyrus and middle temporal regions. The whole-brain analysis for the negative parametric modulation showed significant activations in a set of regions that included the cingulate, precuneus, and striatum.

#### 2) Online activity mediates the cingulo-opercular – memory link

As in Experiment 1, we tested whether activity in regions found in the positive parametric contrast and activity in regions found in the negative parametric contrast mediate the cingulo-opercular - memory link. Both analyses replicated findings of Experiment 1 showing partial mediation, suggesting that around 45% of the link between prestimulus cingulo-opercular activity and successful memory performance was mediated by an increase in clip-related activity in a set of temporal regions (Figure 3c) and by decrease in clip-related activity in a set of regions that included the precuneus, striatum, and cingulate cortex (Figure 3d). Path coefficients for the model that used activity from regions revealed in the positive parametric contrast (+1) as mediator were: path c: b = 0.98, [0.67, 1.3], path a: b = 0.16 [0.14, 0.19], path b: b = 2.67 [1.94, 3.43], path ab: b = 0.44 [0.31, 0.58], path c’: b = 0.54 [0.26, 0.84]. Path coefficients for the model that used activity from regions revealed in the negative parametric contrast (−1) as mediator were: path c: b = 0.93, [0.63, 1.25], path a: b = −0.26 [−0.28 −0.23], path b: b = −1.58 [-2.26, - 0.93], path ab: b = 0.4 [0.23, 0.59], path c’: b = 0.53 [0.23, 0.84]. Thus, the statistical criteria for partial mediation were met in both models, replicating the findings of Experiment 1.

### Linking prestimulus and poststimulus activity

Correlating memory-predictive prestimulus activity (cingulo-opercular regions) with memory-predictive post-clip activity (including hippocampus, striatum, caudate) did not reveal a link between these two time-windows (see Supplementary Information for more details).

## General Discussion

The current study is to the best of our knowledge the first to explore prestimulus brain activity that predicts encoding of novel, naturalistic stimuli. Furthermore, the current study provides a mechanistic account linking the observed prestimulus activity to memory formation via modulation of online stimulus activity. In two independent data sets, we found that prestimulus activity in the cingulo-opercular network correlates with subsequent memory performance. Mediation analyses revealed that prestimulus cingulo-opercular activity gates memory performance by enhancing clip-related activity in temporal regions and by dampening clip-related activity in a set of regions that include the precuneus, cingulate and striatum.

According to the dual model network of attentional control (Dosenbach et al. 2008), the cingulo-opercular network is associated with adaptive control of attention and the initiation and maintenance of task goals while the frontoparietal network is involved in adjustment of control in response to feedback. In the current work memory-predictive prestimulus activations were observed mainly in the cingulo-opercular network, with some involvement of regions considered to be part of the frontoparietal (sometimes referred to as dorsal attention) network. These regions (left precuneus and left inferior parietal) were found to take part in the initiation of task set (Dosenbach et al., 2006). The involvement of these regions, alongside the cingulo-opercular network, suggests that attentional states mainly related to the initiation of task goals preceding an event, play a role in shaping long-term memory. This idea raises a question regarding the nature of the observed memory-predictive attentional state, and specifically whether memory-predictive activity in the cingulo-opercular network results from a deliberate preparatory process or from an incidental attentional state. Several findings of the current work suggest that memory-predictive activation in the cingulo-opercular network was less related to intentional preparation: 1) We did not observe a main effect for instruction type (remember > look) in the cingulo-opercular network in Experiment 1 (see SI3); 2) we did not observe a difference in results of the mediation models for “remember” and “look” trials (see SI10); and 3) we replicated the finding that cingulo-opercular activity predicts memory performance in Experiment 2, in which there was no instruction cue prior to the memoranda. Therefore, we postulate that incidental brain fluctuations in the cingulo-opercular network modulate encoding (see also Turk-Browne et al. 2006; Addante et al. 2015). Specifically, events starting during incidental high activity in this network may be remembered better than events starting during incidental low activity. In support of this notion are imaging (Yoo et al. 2012), electrophysiological (Griffin et al. 2004; Schurger et al. 2012) and intracranial brain stimulation (Ezzyat et al. 2017) findings showing that prestimulus brain oscillations can influence memory-related processes.

While the current work focused on cingulo-opercular activity in the prestimulus phase, an examination of the time course reveals that this network plays an opposite role during stimulus presentation. Specifically, during the clips, the cingulo-opercular network was deactivated more strongly for subsequently-remembered clips compared to subsequently forgotten ones. This is in line with previous findings (e.g., Daselaar et al. 2004), and may suggest that processes needed for the preparation of efficient encoding during the prestimulus phase are no longer needed (and should even be suppressed) during the event. This finding may also help reconcile the mixed findings in the literature regarding the memory-predictive effect of cingulo-opercular activity during stimulus presentation (e.g., Vaden et al. 2017).

Our findings suggest both direct and indirect influence of prestimulus cingulo-opercular activity on memory performance. Specifically, using a multi-level logistic mediation analyses we showed that the link between prestimulus cingulo-opercular activity and memory is partially mediated by clip-related activity in two distinct networks. Namely, elevated activity in the cingulo-opercular network was associated with enhanced activity in regions such as the fusiform and middle temporal gyrus, which are thought to play a role in encoding (for a meta-analyses see Spaniol et al. 2009; Murty et al. 2010; Kim 2011), and with reduced activity in a set of regions that were found to be involved in retrieval and self-referential processing (for meta-analysis and review papers see: Wagner et al. 2005; Northoff et al. 2006; Spaniol et al. 2009; Kim et al. 2010). These results support previous findings showing a competitive relationship between networks involved in encoding and retrieval (Kim et al. 2010; Kuhl et al. 2011) and suggest a gating role for attention in determining which of these processes will take precedence. Specifically, as attention plays a prominent role in shifting between external and internal focus (Chun et al. 2011; Kucyi et al. 2017), it is possible that prestimulus attentional state enhances encoding by promoting external focus as well as by suppressing interference by internally-generated thoughts (e.g., retrieval of past memories). This suggestion is speculative and should be tested in studies which directly manipulate retrieval/internal focus.

Previous findings suggest involvement of the hippocampus in both prestimulus (Park and Rugg 2010; Addante et al. 2015) and poststimulus (Ben-Yakov and Dudai 2011a; Ben-Yakov et al. 2013, 2014) phases. In the current work we did not observe memory-predictive prestimulus activity in the hippocampus. The reason for the difference between our findings and findings of prior work (Park and Rugg 2010; Addante et al. 2015) may be related to the memory probed. Specifically, in the current work we tested participants’ memory for the gist of naturalistic events while other studies tested recollection of pictures or words. This suggestion, however, is hypothetical and should be addressed in future work. We did detect, in line with previous findings (Ben-Yakov and Dudai 2011a; Ben-Yakov et al. 2013, 2014), memory-predictive hippocampal activity at clip offset. Interestingly, memory-predictive prestimulus activity (cingulo-opercular regions) was not correlated with memory-predictive post-clip activity (including hippocampus, striatum, caudate). These finding, although yet speculative, suggests that the processes underlying the role of prestimulus activity in encoding are different from those linking post-event activity to memory success.

Most previous fMRI studies that explored the association between prestimulus activity and memory performance presented a cue that predicted the content of the to-be-remembered event (Adcock et al. 2006; Mackiewicz et al. 2006; Wittmann et al. 2007; Park and Rugg 2010; Addante et al. 2015), raising the possibility that the anticipation for specific content modulated the observed effects. In the current study, the results cannot be explained by anticipation because no relevant cue was given prior to the movie clips and the effects of degree of temporal anticipation were accounted for in the design (Experiment 1) and analysis (Experiments 1 & 2; see SI). Additional control analyses (reported in the SI) ruled out sequential effects (i.e., effects related to memory performance in the previous clip), as well as arousal influences (as indicated by a parametric analysis that included pupil dilation). Furthermore, we demonstrated that findings of the current work cannot be explained by temporal anticipation or by overlap between prestimulus and stimulus-related activity. Therefore, the findings of the current work lead to several predictions that may be tested in further studies. Specifically, real-time fMRI and TMS/tDCS/intracranial stimulation studies can provide direct evidence for the role of spontaneous cingulo-opercular fluctuations in memory success. Furthermore, studies manipulating attention and task-goals can provide evidence for the role on intentional attentional states in enhancing encoding and in reducing interference by internal focus. These studies can also shed light on the specificity of the findings to the cingulo-opercular network and the role of frontoparital regions in the observed memory-predictive prestimulus effect.

In conclusion, we propose that prestimulus attentional states as reflected in cingulo-opercular activity may enhance memory encoding by shifting the balance between encoding and retrieval – increasing focus on the external environment while reducing interference from task-unrelated, internally generated, memories.

## Acknowledgments

The authors declare that they had no conflicts of interest with respect to their authorship or the publication of this article. We wish to thank Sam Hutton for his help with the programming of Experiment 1, Yael Holtzman for her help in running participants, Matti Vuorre for his help with the multi-level mediation analysis, Ronen Hershman for his help with the pupillometry analysis, and Edna Haran and the Weizmann MRI technician team, for assistance in the imaging scans. We also wish to thank Daphna Shohamy and Mariam Aly for helpful ideas and suggestions during analysis and manuscript preparation.

