## Supplementary for "Prestimulus Activity in the Cingulo-Opercular Network Predicts Memory for Naturalistic Episodic Experience"

**Data and Code**

For data sharing purposes, all data have been organized according to the Brain Imaging Data Structure (BIDS) standards (Gorgolewski et al, 2016), and are freely available via the OpenNeuro repository (https://openneuro.org; Prestimulus Exp1 and Prestimulus Exp2; Poldrack and Gorgolewski, 2017). T-maps and scripts are available via the OSF website (https://osf.io/x3q2r/). Note that the conversion to BIDS was done post-hoc, after manuscript preparation, so the original scripts should be run with after adaptation to the BIDS data.

**Experiment 1**

**Inter-rater reliability of the recall scoring**

The reliability of the scoring (remembered/forgotten/unclear) was assessed post-hoc (after the analysis had already been run) by asking two naïve raters to label the clips. This analysis revealed high similarity between the original scoring and that of each naïve rater (Cohen's Kappa rater 1 = .843, 95% CI, .826 to .859, p < .0005; Cohen's Kappa rater 2 = .828, 95% CI, .811 to .845, p < .0005).

**Eye-tracking Data**

An MR-compatible eye tracker (EyeLink1000, SR Research, Ontario, Canada) with a sampling rate of 500 Hz (2 ms inter-sampling time) was used to collect eye-movements, blinks, and pupil size during the Study session. Stimulus presentation and data acquisition were controlled by EyeLink software (Experiment Builder). Participants were requested to keep their eyes fixated on the center of the screen and to avoid eye movements for the entire task. Pupil area was determined using EyeLink's “centroid” algorithm. Two participants were excluded from analysis due to a large proportion of missing samples (over 50% points in time when the eye-tracking software did not identify the pupil) leading to omission of more than 25% of their trials (resultant sample: N = 19, 6 males, mean age = 25.5 ± 3.6).

***Average fixation duration, number of fixations, number of blinks, or number of saccades.***

Repeated-measures ANOVAs were used to test the main effect of instruction type (remember/look), the main effect of memory (remembered/forgotten) and the interaction between them on several dependent variables: fixation duration, number of fixations, number of missing values (mostly associated with blinks), and number of saccades. These dependent variables were extracted from DataViewer, an analysis tool provided by Eye-link, and included only samples within the prestimulus (instruction) time window. There were no significant main effects or interaction for any of these measures (all p’s > .20).

***Pupil size***

Pupil size is a physiological marker of arousal and cognitive load (Kahneman and Beatty 1966; Samuels and Szabadi 2008). As described in the manuscript, a parametric analysis was performed in order to examine whether processes related to arousal /cognitive load may explain our neuroimaging findings.

***Pupillometry data pre-processing***

Pupil data was processed in MATLAB R2016a (MathWorks) using CHAP (Hershman et al. 2019) in a procedure similar to that used in previous studies (e.g., Cohen et al. 2015; Hershman et al. 2018). First, pupil data extracted from the Eye-Link (pupil size in pixels) was z-scored. Z-scoring was done for each participant separately, on the entire time course (i.e., based on the mean and SD calculated on all trials). Linear interpolation was used to fill missing values. Trials with over 50% missing pupil values (Mean = 10.2%, SD = 5.7%) were excluded from the analysis. For each trial, as the fixation at the beginning of the trial varied in duration (due to opening and closing of the camera), a 200-ms pre-instruction baseline average was subtracted from each of the following pupil samples. This baseline was used to eliminate potential influences from the pre-instruction period. Per-trial prestimulus pupil size was calculated as the average pupil size in the instruction time-window (from instruction’s onset to clip’s onset).

For illustration purposes, we present time courses during the instruction and clip periods for remember vs forgotten clips and for look vs instruction condition when no baseline correction was used. As seen in the figure, there was no difference in pupil size between the memory and instruction conditions during both instruction and clip time-windows.

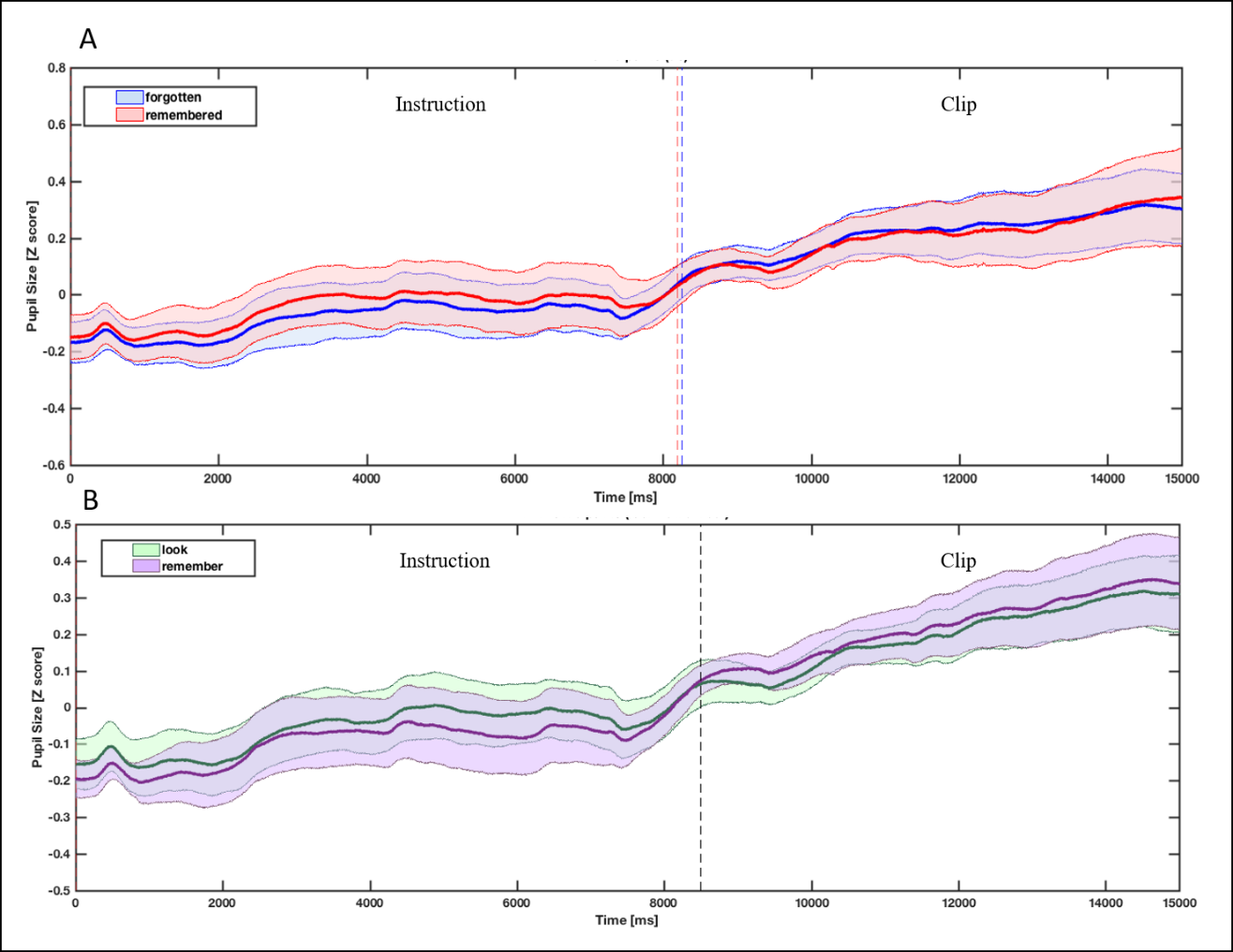

**SI1.** Pupil size time-course for the A) memory condition (remembered vs forgotten) and B) instruction type (“Remember”/”Look). Dashed lines represent the mean offset of the instruction cue. Error bars denote 95% CI.

**fMRI**

**Whole-brain results for the remembered > forgotten contrast**

| Side | Region | MNI Coordinates  (x, y, z) | | | t-value | Voxels |
| --- | --- | --- | --- | --- | --- | --- |
| L | Anterior Prefrontal | -27 | 38 | 7 | 6.36 | 375 |
| R | Anterior Prefrontal + Insula | 33 | 47 | 22 | 6.12 | 234 |
| R | Superior Frontal | 15 | 62 | 1 | 4.99 | 232 |
| L | Anterior Insula | -33 | 14 | 4 | 4.65 | 53 |
| R | Postcentral | 36 | -40 | 58 | 4.59 | 85 |
| L | Precuneus | 12 | -70 | 43 | 4.41 | 126 |
| R | Dorsal Anterior Cingulate | 3 | 20 | 37 | 4.36 | 175 |
| L | Inferior Parietal | -36 | -52 | 40 | 4.28 | 62 |

### *SI2.* Brain activity for the whole-brain analysis of the remembered > forgotten contrast (p < .001, cluster pFWE < .05). Note that the right anterior prefrontal cluster extended to the right anterior insula.

**Whole-brain results for the remember > look contrast**

The contrast “remember” vs “look” revealed bilateral occipital activity (see Table SI2).

| Side | Region | MNI Coordinates  (x, y, z) | | | t-value | Voxels |
| --- | --- | --- | --- | --- | --- | --- |
| L | Occipital | -33 | -100 | -8 | 6.63 | 220 |
| R | Occipital | 39 | -94 | -5 | 5.69 | 87 |

### *SI3.* Peak coordinates and cluster size for the remember > look contrast (p < .001, cluster pFWE < .05).

**Time-course illustration**

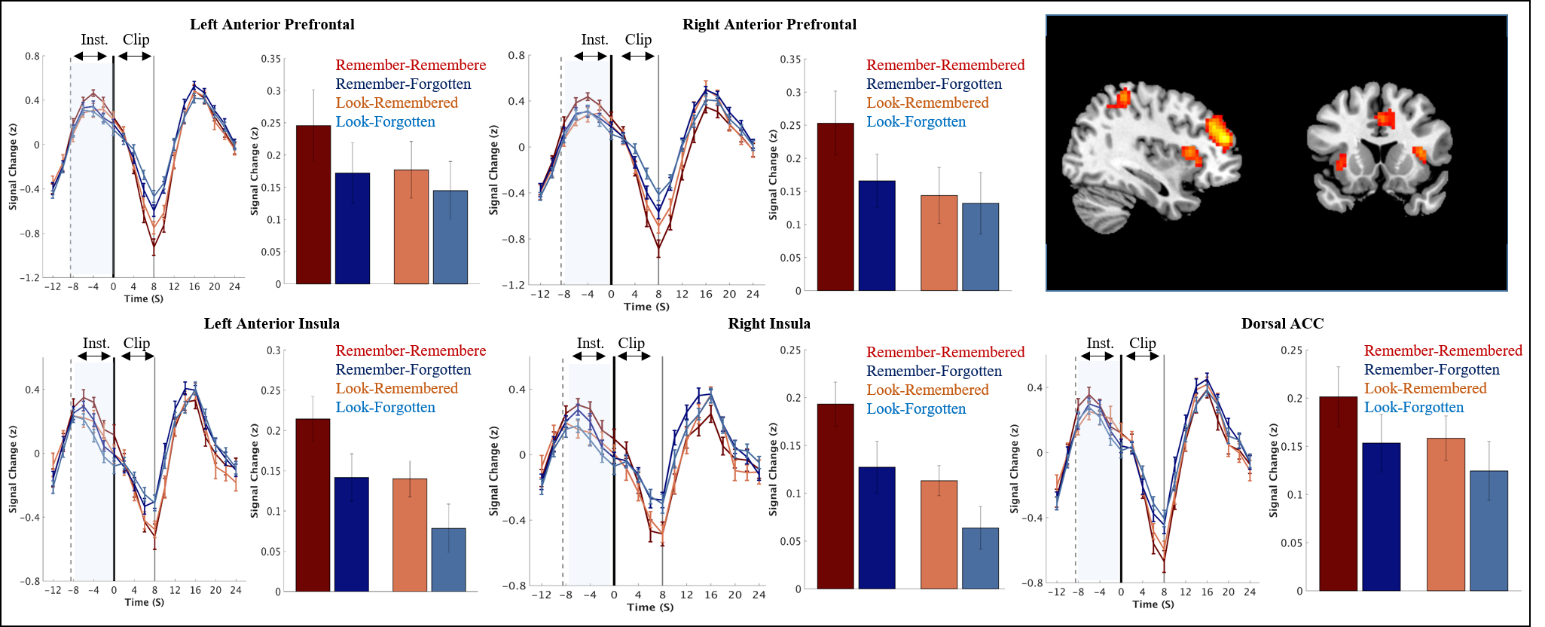

***SI4****.* For illustration purposes, mean group BOLD signal (after z scoring each time course) were extracted from regions of the cingulo-opercular network using a functional ROI. The black lines indicate the onset of clip presentation, the gray lines indicate the offset of the current clips, and the dashed lines represent the mean onset of the instruction cue. The bar figures represent the mean activity during the instruction time-window for each of the conditions.

**Controlling for anticipation, clip-related activity, and arousal**

We computed the statistics for three additional first-level models. All three models showed nearly identical results as the original model.

1) A model that included the duration of the instruction period as a parameter was used to control for anticipatory influences. Temporal anticipation is high when interval lengths are fixed, but is also present in protocols using fixation/rest periods of varying lengths (Turk-Browne et al., 2006), a phenomenon called the foreperiod effect (Niemi & Näätänen, 1981). For example, in event-related (compared to block) designs, longer intervals are associated with increased anticipation (Niemi and Näätänen 1981). Thus, by controlling for instruction length we were able to rule out remembered > forgotten effects driven by temporal anticipation (see SI4(. Furthermore, cingulo-opercular activity showed a similar pattern across the different instruction durations (see SI5) and looking at the parametric analysis in which instruction duration was entered as a parameter did not reveal activations that are dependent on this parameter (p < .001, cluster pFWE < .05).

| Side | Region | MNI Coordinates  (x, y, z) | | | t-value | Voxels |
| --- | --- | --- | --- | --- | --- | --- |
| L | Anterior Frontal | -27 | 38 | 7 | 6.65 | 395 |
| R | Anterior Frontal + Insula | 33 | 47 | 22 | 5.96 | 229 |
| R | Superior Frontal | 15 | 62 | 1 | 5.14 | 252 |
| R | Postcentral | 36 | -40 | 58 | 4.48 | 65 |
| R | Precuneus | 15 | -70 | 43 | 4.43 | 123 |
| R | Dorsal anterior Cingulate | 3 | 20 | 37 | 4.28 | 179 |
| L | Inferior Parietal | -36 | -52 | 40 | 4.20 | 61 |

### *SI5.* Brain activity for the whole-brain analysis of the remembered > forgotten contrast (p < .001, cluster pFWE < .05) in a model that included instruction duration as a parametric regressor.

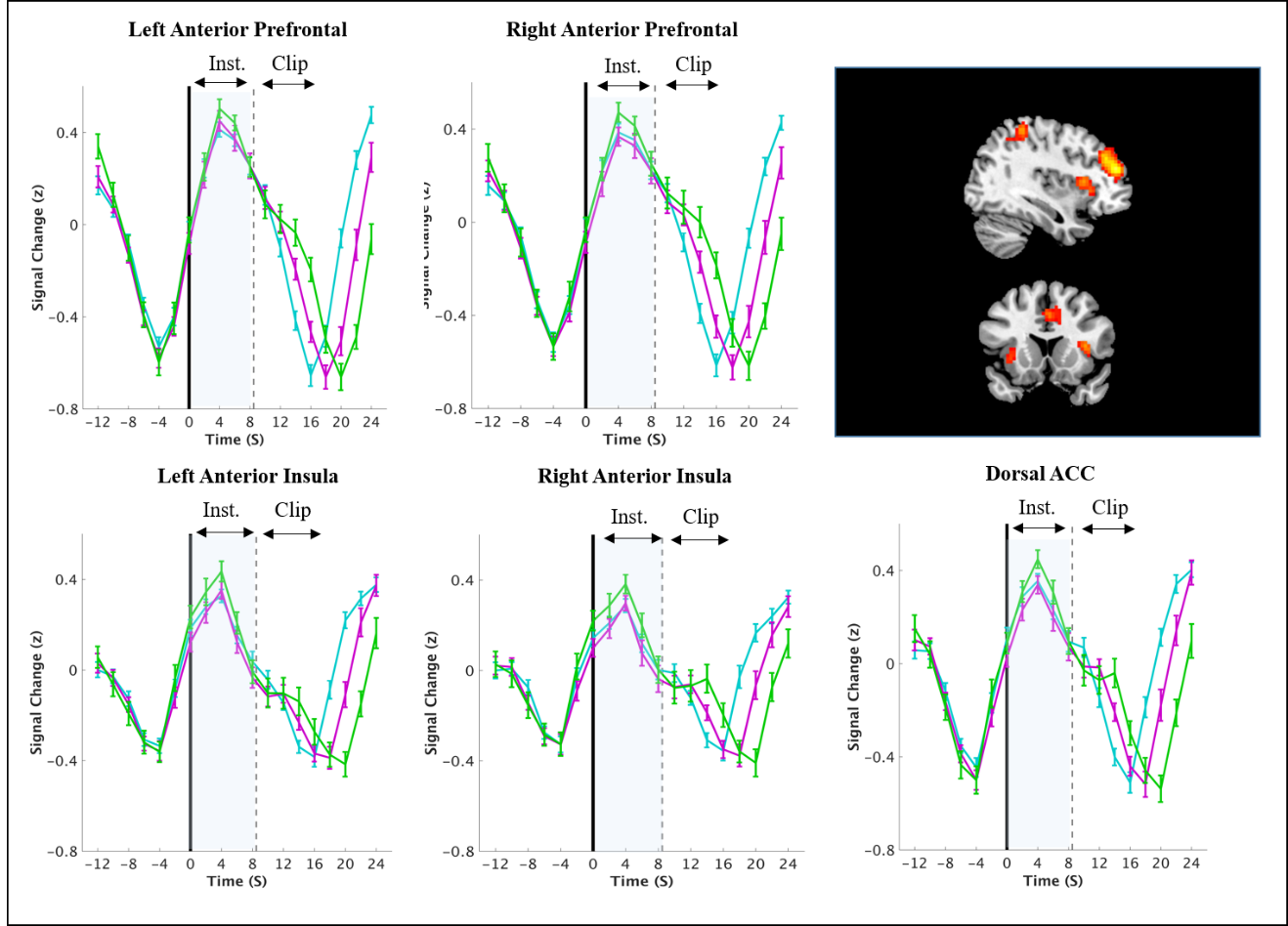

***SI6****.* For illustration purposes, mean group BOLD signal (after z scoring each time course) for each instruction duration were extracted from regions of the cingulo-opercular network using a functional ROI. The black lines indicate the onset of instruction cue (“remember”/ “look”), and the dashed lines represent the mean offset of the instruction cue. Cyan lines represent instruction duration of 7 seconds, magenta lines represent instruction duration of 9 seconds, and green lines represent instruction duration of 11 seconds.

2) Since the focus of the current study was on prestimulus activity, there was no theoretical point in adding a jittered interval between the prestimulus and stimulus phases. The use of long intervals for these two phases (7-11 sec for the prestimulus phase and 8 sec for the clip stimulus) should have eliminated effects resulting from activity overlap between these two phases. However, to make sure that the observed prestimulus effects did not result from clip-related activity, we conducted a model that included both the prestimulus (instruction) and stimulus (clip) time-windows as regressors to prevent systematic variability associated with the actual clip period to influence the prestimulus period beta estimate.

| 3) Side | Region | MNI Coordinates  (x, y, z) | | | t-value | Voxels |
| --- | --- | --- | --- | --- | --- | --- |
| R | Inferior Parietal | 36 | -43 | 58 | 5.60 | 120 |
| L | Anterior Frontal | -27 | 38 | 7 | 5.52 | 207 |
| R | Anterior Frontal | 33 | 47 | 22 | 5.22 | 184 |
| R | Superior Frontal | 30 | -4 | 58 | 4.66 | 73 |
| R | Insula | 33 | 20 | 13 | 4.59 | 110 |
| L | Insula | -33 | 14 | 4 | 4.22 | 68 |
| R | Dorsal Anterior Cingulate | 3 | 20 | 40 | 4.17 | 100 |
| L | Inferior Parietal | -54 | -28 | 43 | 4.11 | 56 |

### *SI7.* Brain activity for the whole-brain analysis of the remembered > forgotten contrast (p < .001, cluster pFWE < .05) in a model that included both prestimulus and clip regressors.

3) A third model that included averaged pupil size during the instruction time window as a parameter was used to control for arousal (Murphy et al., 2014).

| 4) Side | Region | MNI Coordinates  (x, y, z) | | | t-value | Voxels |
| --- | --- | --- | --- | --- | --- | --- |
| L | Anterior Frontal | -27 | 38 | 7 | 6.07 | 249 |
| R | Anterior Frontal + Insula | 36 | 47 | 22 | 5.58 | 196 |
| L | Precuneus | -9 | -70 | 43 | 5.28 | 232 |
|  | Posterior Cingulate | 0 | -31 | 28 | 5.14 | 105 |
| R | Superior Frontal | 15 | 59 | 1 | 4.86 | 154 |
| R | Superior Frontal | 30 | -4 | 58 | 4.77 | 60 |
| L | Inferior Parietal | -36 | -52 | 40 | 4.10 | 60 |
| R | Dorsal Anterior Cingulate | 3 | 17 | 40 | 4.09 | 75 |

### *SI8.* Brain activity for the whole-brain analysis of the remembered > forgotten contrast (p < .001, cluster pFWE < .05) in a model that included prestimulus pupil size as a parametric regressor.

**Controlling for sequential effects**

Two analyses were conducted to test whether memory performance (remembered/forgotten) and instruction type (remember/look) in the previous trial affected our results for the remembered>forgotten contrast. For this purpose, an additional regression model was estimated, in which we constructed separate regressors for each trial. In this single-trial model, we modeled each event during the prestimulus-instruction phase (total of 160 regressors), each convolved with the canonical HRF. For each of the prestimulus periods, we then extracted the ROI-averaged beta (amplitude) estimates for the brain regions found in the remembered > forgotten contrast (voxel p < 0.001, cluster pFWE < 0.05). The extracted and averaged betas were then, together with the memory performance and instruction type in the current and previous trial, subjected to two repeated-measures ANOVAs (one assessing the interaction between previous memory performance and current memory performance and a second one assessing the interaction between previous instruction type and current memory performance). These ANOVAs were conducted using IBM SPSS Statistics Version 23.0 (IBM Corp. Released 2015. IBM SPSS Statistics for Windows, Version 23.0. Armonk, NY: IBM Corp.)

We found no indication for sequential effects. Specifically, there was no main effect for previous trial’s memory performance (F(1,21) = 1.21, p = 0.28) or instruction type (F(1,21) < 1), and no interaction between current memory performance and previous trial’s memory performance (F(1,21) = 2.12, p = 0.16) or instruction type (F(1,21) < 1).

**Brain activity for the whole-brain parametric analysis of the positive (+1) and negative (-1) contrasts**

| Side | Region | MNI Coordinates  (x, y, z) | | t-value | | Voxels | |
| --- | --- | --- | --- | --- | --- | --- | --- |
| Positive Parametric Contrast (+1) | | | | | | | |
| R | Inferior Frontal | 57 | 32 | 13 | 9.61 | | 110 |
| R | Amygdala/Hippocampus | 24 | -7 | -8 | 7.71 | | 129 |
| L | Amygdala/Hippocampus | -24 | -10 | -11 | 7.27 | | 144 |
| R | Superior Frontal | 9 | 53 | 40 | 6.51 | | 132 |
| R | Fusiform | 24 | -40 | -11 | 5.82 | | 164 |
| L | Fusiform | -33 | -58 | -8 | 5.67 | | 97 |
| R | Middle Occipital | 42 | -85 | 22 | 5.53 | | 141 |
| L | Middle Temporal | -48 | -28 | 1 | 5.52 | | 211 |
| R | Middle Temporal | 57 | -7 | -11 | 5.16 | | 193 |
| Negative Parametric Contrast (-1) | | | | | | | |
| R | Middle Cingulate (extending to Striatum; Putamen, Caudate) | -3 | -25 | 37 | 6.17 | | 2434 |
| R | Precuneus | 12 | -67 | 37 | 8.87 | | 1332 |
| L | Inferior Parietal | -36 | -55 | 43 | 7.92 | | 460 |
| R | Inferior Parietal | 48 | -55 | 43 | 7.92 | | 311 |
| R | Middle Temporal | 36 | -43 | 1 | 6.56 | | 112 |
| R | Middle Frontal | 36 | 44 | 16 | 5.07 | | 105 |

### *Table SI9.* Brain activity for the whole-brain parametric analysis of the positive (+1) and negative (-1) contrasts in Experiment 1. Prestimulus cingulo-opercular activity served as a parameter to predict correlated clip-related activity (p < .001, cluster pFWE < .05).

**Mediation for “Remember” and “Look” trials**

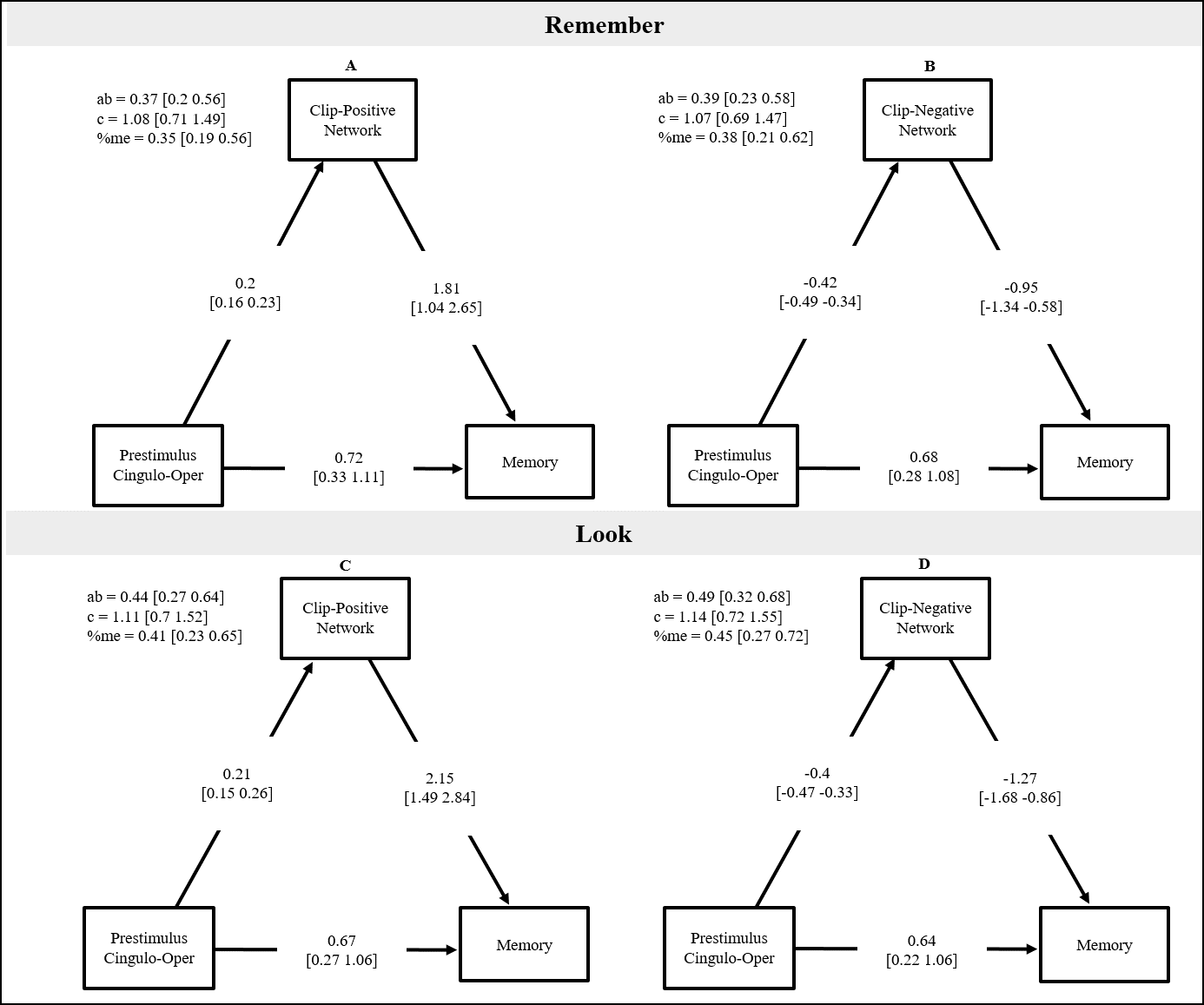

***SI10.*** Multi-level mediation model for remember and look trials.

**Experiment 2**

**Inter-rater reliability of the recall scoring**

The reliability of the scoring (remembered/forgotten/unclear) was assessed post-hoc by asking two naïve raters to label the clips. This analysis revealed high similarity between the original scoring and that of each naïve rater (Cohen's Kappa rater 1 = 714, 95% CI, .686 to .740, p < .0005, Cohen's Kappa rater 2 = .826, 95% CI, .801 to .849, p < .0005).

**fMRI**

**Coordinates of the cingulo-opercular network**

| Side | Region | MNI Coordinates  (x, y, z) | | |
| --- | --- | --- | --- | --- |
| L | Anterior Prefrontal | -28 | 51 | 15 |
| R | Anterior Prefrontal | 27 | 50 | 23 |
| L | Anterior Insula | -35 | 14 | 5 |
| R | Anterior Insula | 36 | 16 | 4 |
|  | Dorsal Anterior Cingulate | -1 | 10 | 46 |

***Table SI11.*** Coordinates of the cingulo-opercular network based on Dosenbach et al (2007).

**SVC analysis for the remembered > forgotten contrast**

| Side | Region | MNI Coordinates  (x, y, z) | | | t-value | Voxels |
| --- | --- | --- | --- | --- | --- | --- |
| L | Anterior Prefrontal | -33 | 41 | 13 | 6.56 | 58 |
|  | Dorsal Anterior Cingulate | 0 | 11 | 43 | 6.33 | 86 |
| R | Anterior Prefrontal | 36 | 50 | 22 | 5.82 | 44 |
| L | Anterior Insula | -36 | 11 | -2 | 4.78 | 25 |
| R | Anterior Insula | 39 | 14 | -2 | 4.32 | 14 |

### *Table SI12.* Brain activity for the SVC analysis testing for cingulo-opercular activity in the remembered > forgotten contrast during the prestimulus time-window (p < .001, cluster pFWE < .05).

**Whole-brain results for the remembered > forgotten contrast**

| Side | Region | MNI Coordinates  (x, y, z) | | | t-value | Voxels |
| --- | --- | --- | --- | --- | --- | --- |
| R | Superior Frontal/Cingulate | 6 | 29 | 37 | 5.84 | 286 |
| R | Anterior Prefrontal | 39 | 50 | 19 | 5.56 | 458 |
| L | Anterior Prefrontal | -33 | 56 | 16 | 5.43 | 166 |
| R | Inferior Parietal | 51 | -46 | 55 | 4.93 | 253 |
| L | Anterior Insula | -33 | 14 | -2 | 4.34 | 54 |
| R | Posterior Cingulate | 6 | -22 | 25 | 4.30 | 109 |
| L | Inferior Parietal | -39 | -49 | 40 | 4.19 | 87 |

***SI13***. Brain activity for the whole-brain analysis of the remembered > forgotten contrast (p < 0.001, cluster pFWE < 0.05).

**Controlling for anticipation and clip-related activity**

Two additional first-level analyses we conducted, one that included the duration of the prestimulus period as a parameter (see SI6), and a second model that included the clip as an additional regressor (see SI7). Both models showed similar results to the original model. The model in which both prestimulus and clip periods were entered as regressors revealed less activations, as this model includes very little time that is not modeled, which increases the error of the parameter estimates.

| Side | Region | MNI Coordinates  (x, y, z) | | | t-value | Voxels |
| --- | --- | --- | --- | --- | --- | --- |
| R | Dorsal Anterior Cingulate | 6 | 26 | 40 | 5.88 | 339 |
| R | Anterior Frontal | 39 | 50 | 22 | 5.75 | 478 |
| L | Anterior Frontal | -33 | 56 | 16 | 5.40 | 172 |
| R | Inferior Parietal | 51 | -46 | 55 | 4.91 | 254 |
| R | Posterior Cingulate | 6 | -22 | 25 | 4.34 | 115 |
| L | Inferior Parietal | -45 | -49 | 49 | 4.21 | 109 |

***SI14***. Brain activity for the whole-brain analysis of the remembered > forgotten contrast (p < 0.001, cluster pFWE < 0.05) in a model that included the prestimulus period as a parametric regressor.

| Side | Region | MNI Coordinates  (x, y, z) | | | t-value | Voxels |
| --- | --- | --- | --- | --- | --- | --- |
| R | Dorsal Anterior Cingulate | 6 | 29 | 34 | 4.19 | 60 |
| R | Anterior Frontal | 57 | 47 | 1 | 4.11 | 51 |
| L | Inferior Parietal | 48 | -46 | 58 | 3.99 | 54 |

***SI15***. Brain activity for the whole-brain analysis of the remembered > forgotten contrast (p < 0.001, cluster pFWE < 0.05) in a model that included also clip period as a regressor.

**Brain activity for the whole-brain parametric analysis of the positive (+1) and negative (-1) contrasts**

| Side | Region | MNI Coordinates  (x, y, z) | | | t-value | | Voxels | |
| --- | --- | --- | --- | --- | --- | --- | --- | --- |
| Positive Parametric Contrast (+1) | | | | | | | | |
| R | Fusiform & Middle Temporal Gyrus | 30 | -67 | -8 | | 9.06 | | 1572 |
| L | Fusiform & Middle Temporal Gyrus | -30 | -61 | -5 | | 7.92 | | 901 |
| R | Precuneus | 15 | -46 | 46 | | 6.48 | | 138 |
| R | Inferior Frontal | 57 | 32 | 1 | | 5.88 | | 189 |
| L | Cerebellum | -12 | -70 | -38 | | 5.87 | | 97 |
|  | Rectus | 0 | 56 | -17 | | 5.24 | | 53 |
| Negative Parametric Contrast (-1) | | | | | | | | |
| L | Middle Cingulate | -3 | -23 | 31 | | 13.73 | | 1009 |
| R | Middle Frontal | 30 | 50 | 19 | | 10.85 | | 360 |
| L | Middle Frontal | -36 | 47 | 16 | | 10.05 | | 769 |
| L | Precuneus | -9 | -67 | 46 | | 8.33 | | 370 |
| L | Middle Cingulate | -3 | -22 | 31 | | 7.90 | | 176 |
| L | Insula | -36 | 17 | 4 | | 7.36 | | 475 |
| R | Striatum (Caudate, Putamen) | 18 | 20 | -5 | | 6.81 | | 145 |
| R | Insula | 33 | 23 | 7 | | 6.73 | | 222 |
| L | Inferior Parietal | -39 | -49 | 43 | | 6.72 | | 283 |
| L | Striatum (Caudate, Putamen) | -15 | 17 | -5 | | 6.23 | | 47 |
| R | Inferior Parietal | 54 | -43 | 40 | | 5.67 | | 107 |
| R | Middle Cingulate | 6 | -28 | 46 | | 5.57 | | 44 |

### *Table SI16.* Brain activity for the whole-brain parametric analysis of the positive (+1) and negative (-1) contrasts in Experiment 2. Prestimulus cingulo-opercular activity served as a parameter to predict correlated clip-related activity (p < .001, cluster pFWE < .05).

**Does prestimulus activity predicts post-clip activity?**

Prior work found that activity at clip offset, in hippocampus and other regions predicts memory performance (for review see Cohen, Pell, et al. 2015). In order to test whether cingulo-opercular prestimulus activity is correlated with post-clip activity we extracted betas for each trial that consisted a narrative clip during these two time-windows. Betas were extracted from masks based on the wholebrain contrast remember > forgotten (cingulo—opercular regions for the prestimulus phase; hippocampus, caudate, striatum for the post-clip phase). We then ran a multi-level regression model to test whether prestimulus activity predicts post-clip activity. This analysis did not indicate a link between prestimulus and post-clip activity, b = -0.02 [-0.06 0.02]. Therefore, it seems that the mechanism driving prestimulus effect on memory is different than the mechanism involved in post-clip effect on memory. See Figure SI12 for data of individual subjects.

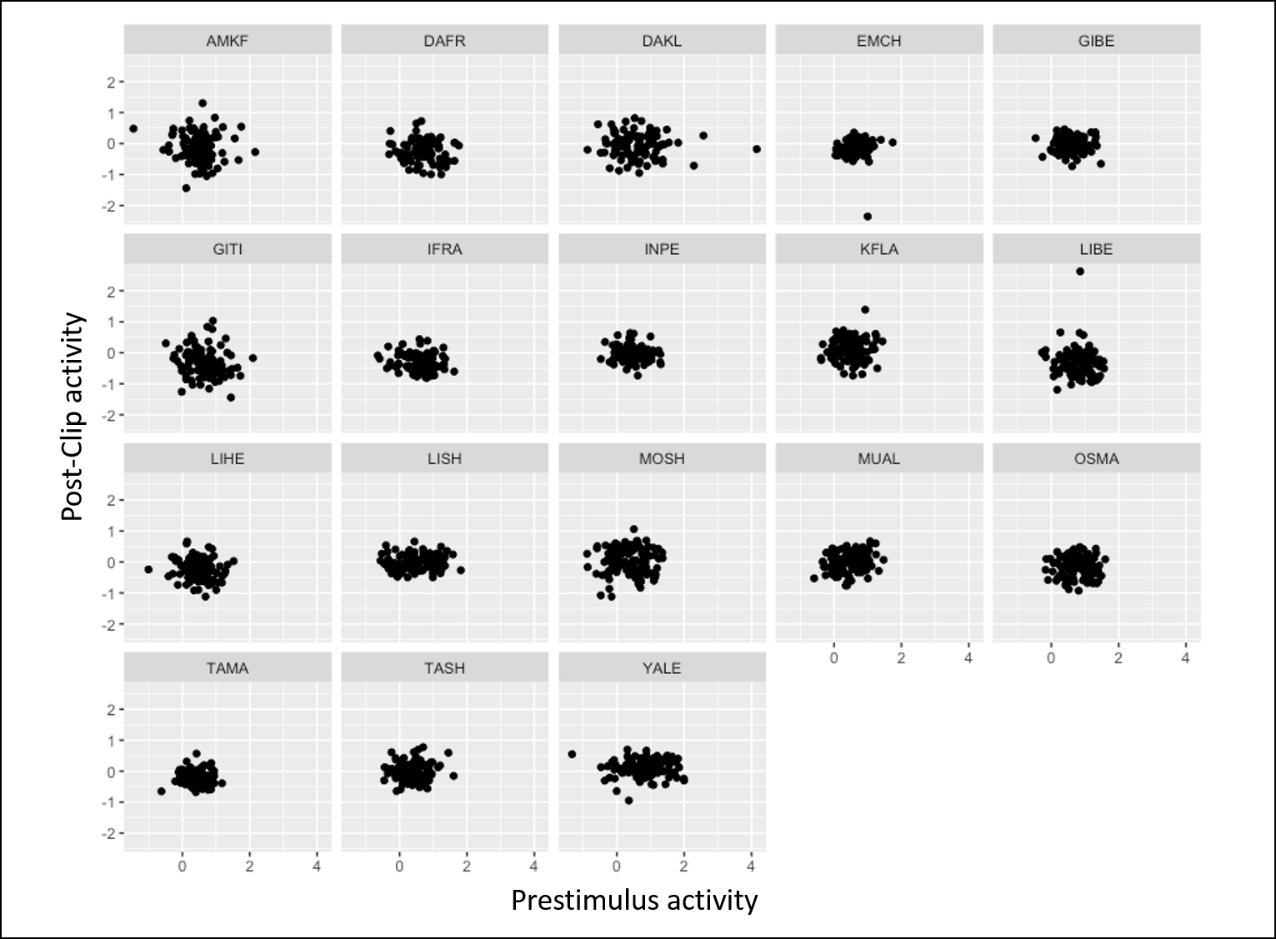

***SI17***. Individual data for prestimulus and post-clip activity. Values on X and Y axes are beta weights (centered).
